# Pharmacological HDAC3 inhibition alters memory updating in young and old mice

**DOI:** 10.1101/2024.05.08.593015

**Authors:** Chad W. Smies, Lauren Bellfy, Destiny S. Wright, Sofia S. Bennetts, Mark W. Urban, Chad A. Brunswick, Guanhua Shu, Janine L. Kwapis

## Abstract

Long-term memories are not stored in a stable state but must be flexible and dynamic to maintain relevance in response to new information. Existing memories are thought to be updated through the process of reconsolidation, in which memory retrieval initiates destabilization and updating to incorporate new information. Memory updating is impaired in old age, yet little is known about the mechanisms that go awry. One potential mechanism is the repressive histone deacetylase 3 (HDAC3), which is a powerful negative regulator of memory formation that contributes to age-related impairments in memory formation. Here, we tested whether HDAC3 also contributes to age-related impairments in memory updating using the Objects in Updated Locations (OUL) paradigm. We show that blocking HDAC3 immediately after updating with the pharmacological inhibitor RGFP966 ameliorated age-related impairments in memory updating in 18-m.o. mice. Surprisingly, we found that post-update HDAC3 inhibition in young (3-m.o.) mice had no effect on memory updating but instead impaired memory for the original information, suggesting that the original and updated information may compete for expression at test and HDAC3 helps regulate which information is expressed. To test this idea, we next assessed whether HDAC3 inhibition would improve memory updating in young mice given a weak, subthreshold update. Consistent with our hypothesis, we found that HDAC3 blockade strengthened the subthreshold update without impairing memory for the original information, enabling balanced expression of the original and updated information. Together, this research suggests that HDAC3 may contribute to age-related impairments in memory updating and may regulate the strength of a memory update in young mice, shifting the balance between the original and updated information at test.

## Introduction

The ability to learn is critical for survival across species. While research has heavily explored the mechanisms of initial memory formation, we know far less about how existing memories are modified with new information. The ability to modify or update existing memories, however, is critical to ensure accurate behavioral responses. Further, memory updating is impaired in old age, contributing to age-related cognitive impairments (Barense et al., 2002; Bizon et al., 2012; Kwapis et al., 2020; Schoenbaum et al., 2002). Despite its fundamental importance to survival and everyday functioning, we have yet to understand the molecular processes that support memory updating and we know even less about how these mechanisms change in old age. Understanding the mechanisms supporting memory updating may shed light on potential therapeutics for memory deficits observed in natural aging and psychiatric disorders.

A newly formed memory must first undergo consolidation to stabilize into long-term memory, a process that requires transcription and translation. During consolidation, a memory is initially labile, susceptible to disruption by a number of amnesic agents, including inhibitors of protein or mRNA synthesis (Davis & Squire, 1984; Duvarci et al., 2008; Igaz et al., 2002; Nader et al., 2000). Once consolidated, however, memories are resistant to disruption by these same amnesic agents. A second period of lability can be induced by memory retrieval, initiating a process termed reconsolidation. As with consolidation, memory reconsolidation requires transcription and translation for successful restabilization. Accumulating evidence demonstrates that the retrieval trial must contain some new information to initiate reconsolidation; when only familiar information is presented at retrieval, the existing memory remains stable and is not susceptible to amnesic agents (De Oliveira Alvares et al., 2013; Díaz-Mataix et al., 2013; Jarome et al., 2015; Kwapis, Jarome, et al., 2017; Kwapis et al., 2020; Morris et al., 2006; Sevenster et al., 2012, 2013, 2014). Thus, one purpose of this reconsolidation process may be to allow existing memories to be modified or updated in response to new, relevant information. Age-related disruption of the reconsolidation process could therefore contribute to reduced memory updating and flexibility in old age.

Epigenetic mechanisms are well-positioned to play an important role in memory updating and contribute to age-related impairments in this process. Epigenetic mechanisms, which modify gene expression without changing the DNA sequence itself, dynamically regulate transcription during memory consolidation (Alaghband et al., 2017; Barchiesi et al., 2022; Kwapis, Alaghband et al., 2017; Monsey et al., 2011) and reconsolidation (Da Silva et al., 2008; Gräff et al., 2014; Lattal & Wood, 2013). In particular, histone acetylation is a key epigenetic mechanism that is critical for memory consolidation and reconsolidation (Bannister & Kouzarides, 2011; Bredy & Barad, 2008; Felsenfeld, 2014; Gräff et al., 2014; Lubin et al., 2011; Maddox & Schafe, 2011; Xhemalce et al., 2011). Histone acetylation enables a permissive state that promotes gene expression (Grunstein, 1997; Strahl & Allis, 2000) and enhances memory formation (Peixoto & Abel, 2013; Ramzan et al., 2020; Villain et al., 2016). Histone acetylation is dynamically regulated through the competing actions of two classes of enzymes: histone acetyltransferases (HATs), which promote acetylation, and histone deacetyltransferases (HDACS), which reduce acetylation (Wapenaar & Dekker, 2016; Wolffe, 1996). HDAC3, the most highly expressed Class I HDAC in the brain, is an especially potent negative regulator of memory consolidation across several different types of memory, including fear conditioning (Kwapis, Alaghband et al., 2017), fear extinction (Alaghband et al., 2017), object location memory (Kwapis et al., 2019; Malvaez et al., 2013; McQuown et al., 2011), object recognition memory (McQuown et al., 2011), cocaine-context place preference consolidation (Rogge et al., 2013), auditory reward learning (Bieszczad et al., 2015), and cocaine place preference extinction (Malvaez et al., 2013). Typically, pharmacological or genetic disruption of HDAC3 around the time of learning transforms a weak learning event into one that establishes a strong and lasting memory, suggesting HDAC3 limits memory strength and persistence during consolidation.

Deletion or disruption of HDAC3 in the dorsal hippocampus of old mice is also sufficient to ameliorate age-related impairments in object location memory, indicating HDAC3 also plays a role in age-related impairments in memory consolidation (Kwapis et al., 2018). Although HDAC3 plays a clear role in initial memory consolidation, whether it plays a similar role in reconsolidation-dependent memory updating is currently unknown. Further, it is also unclear whether HDAC3 also contributes to age-related impairments in memory updating.

To better understand the role of HDAC3 in memory updating in both young and old animals, we used our recently developed spatial memory updating paradigm, the objects in updated locations (OUL) task. OUL leverages a mouse’s innate attraction to novelty to determine how a memory update incorporates into an existing object location memory. Importantly, OUL can assess both the original memory and the updated information in a single, non-stressful test session. In addition, OUL depends on the dorsal hippocampus, a brain region critical for memory and affected early in aging (Aziz et al., 2019; Bach et al., 1999; Kwapis et al., 2018). OUL is therefore ideal for understanding both the mechanisms that support memory updating and the mechanisms that contribute to age-related updating deficits. Here, using the pharmacological HDAC3 inhibitor RGFP966 (Bieszczad et al., 2015; Malvaez et al., 2013; Phan et al., 2017; Shang et al., 2019), we tested the role of HDAC3 in memory updating in young and old mice. Our results show that HDAC3 inhibition after memory updating enhanced memory for the update in old mice but, to our surprise, impaired memory for the original information in young mice. A follow-up study then determined that the original and updated information compete for expression, with HDAC3 inhibition shifting which information is expressed at test. Overall, our studies suggest that HDAC3 negatively regulates reconsolidation-dependent memory updating in young and old mice and contributes to memory updating impairments in old mice.

## Methods

### Animals

Adult (3-month-old) male C57BL/6J mice were obtained from Jackson Laboratories (JAX), and old (18-20-month-old) male C57BL/6J mice were obtained from the National Institute on Aging (NIA) Aged Rodent Colony. We specifically chose 18-20-m.o. mice, as this is when we and others have previously observed deficits in memory and similar behaviors (Bach et al., 1999; Brito et al., 2023; Ederer et al., 2022; Hendrickx et al., 2022; Kwapis et al., 2018; Shoji et al., 2016; Weber et al., 2015), including age-related impairments in OUL (Kwapis et al., 2020). All animals were maintained in a temperature (68-79°C) and humidity-controlled (30-70%) environment under a 12-hour light/dark cycle (lights on at 7A, lights off at 7P). Behavior was conducted during the light phase, when we observe peak spatial memory performance (Bellfy et al., 2023; Brunswick et al., 2023; Urban et al., 2021). Mice had access to food and water *ad libitum*. All experiments were conducted according to US National Institutes of Health guidelines for animal care and use and were approved by the Institutional Animal Care and Use Committee of Pennsylvania State University. All animals were group-housed with four animals per cage.

### RGFP966 Preparation

A solution containing 30% HP-ß-CD and 100mM Na-acetate was made in sterile water, brought to a pH of 5.4 with HCl, and sterilized with a 0.22µm filter. The sterilized HP-ß-CD solution was then used to make the 1% DMSO vehicle. Finally, a stock solution of 10mg/mL RGFP966 in DMSO was diluted to 0.1mg/mL with the sterile HP-ß-CD solution for systemic subcutaneous injections of 10mg/kg. We chose this dose as it has been shown to cross the blood-brain barrier at sufficient concentrations (Malvaez et al., 2013) and improves memory in multiple tasks (Bieszczad et al., 2015; Malvaez et al., 2013; Shang et al., 2019).

### Objects in Updated Locations (OUL) Paradigm

The Objects in Updated Location task was conducted similarly to that previously described (Wright et al., 2020). Briefly, animals were handled for four days for two minutes/day and habituated to the context for six consecutive days for five minutes per day, with two handling days overlapping with habituation days. Animals were scruffed and weighed following habituation sessions four days prior to training to get accustomed to being scruffed for injections. Animals were then trained with two identical objects (200mL tall form graduated beakers filled with cement) placed in specific locations (A1 and A2) for 10 minutes, either for one or three days, with 24 hours separating each session. Next, 24 hours later, mice were given an update session, in which one of the objects was moved to an updated location (A3). Animals were allowed to explore for five minutes (1 minute for the subthreshold update experiment) and given subcutaneous injections of either RGFP966 or vehicle immediately after the update session. Finally, 24 hours later, mice were given a retention test to assess memory for both the original and the updated object locations. At test, four objects were presented in specific locations within this familiar context: two objects in the original locations (A1 and A2), one object in the updated location (A3), and one object in a completely novel location (A4) (**FIGS 1-3**).

**Figure 1.**
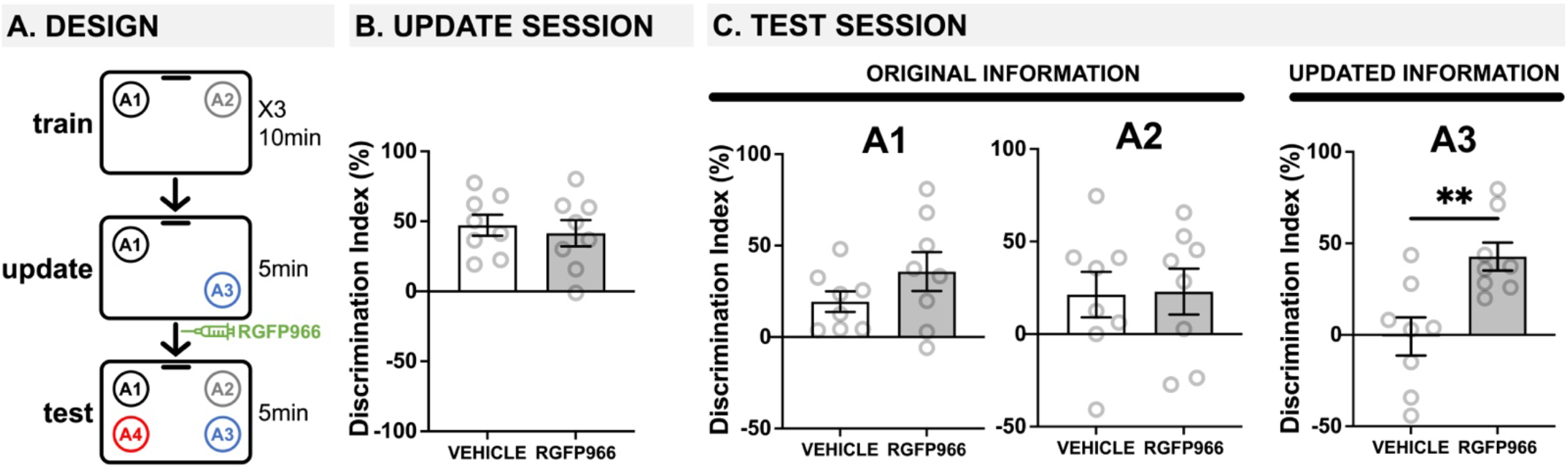
Age-related deficits in memory updating are ameliorated by blocking HDAC3 immediately after updating. (**A**) Experimental design using the OUL paradigm with 18-20-m.o. male mice. Mice were injected with the HDAC3 inhibitor RGFP966 immediately after the update session. (**B**) Update session behavior. No differences were observed during updating, before RGFP966 injection. (**C**) Test session behavior. For locations A1 and A2, no differences were observed between RGFP966-treated and vehicle-treated animals suggesting post-update RGFP966 injections did not impact memory for the original information. For the updated location A3, however, RGFP966 injections improved memory, with RGFP966 mice showing significantly higher DIs for A3 compared to vehicle controls. (**\***p<0.05, **\*\***p<0.01, **\*\*\***p<0.001)

### Data Analysis & Statistics

Behavior videos were collected using Ethovision (Noldus, Leesburg, VA). Habituation data (movement distance and speed) was calculated within the Ethovision software for all behavioral experiments; a reduction in movement and velocity was used to indicate successful habituation to the context (see **FIGS S1-S3**). Behavioral videos were manually scored by experimenters blinded to the experimental groups using Deepethogram (Bohnslav et al., 2021) or Object Task Timer (UC Irvine). Deepethogram and Object Task methods were validated within and between experimenters. The criteria for behavioral scoring followed those described in (Wright et al., 2020). For each session, we calculated a Discrimination Index (DI): DI = (t_novel_ – t_familiar_) / (t_novel_ + t_familiar_) x 100%. For the test session, three different DIs were calculated for objects A1, A2, and A3 (familiar) compared to the novel location A4. When both objects were equally novel or familiar (e.g., training), the ‘novel’ and ‘familiar’ locations were randomly chosen and counterbalanced between conditions. Statistical analyses were conducted using GraphPad Prism (RRID:SCR_002798). Any mouse with a Discrimination Index % (DI) more than two standard deviations away from the mean or with an exploration time of less than two seconds during testing or one second for the subthreshold update session (to ensure some exploration during the shortened session time) were removed from all behavioral analyses. Group differences were analyzed with Student’s *t*-tests or 2-way ANOVAs and Sidak’s multiple comparison *post hoc* tests. For each group, to determine if each DI at test was significantly different from a value of 0, indicating no preference for either location, we used one-sample *t-*tests comparing each group to the hypothetical value of 0. In all experiments, an °-value of 0.05 was required for significance. Error bars in all figures indicate SEM.

## Results

### HDAC3 blockade ameliorates age-related impairments in memory updating

We first aimed to test whether HDAC3 inhibition could ameliorate age-related deficits in memory updating. We previously demonstrated that 18-month-old mice show impaired memory updating in OUL even when the original memory is intact (Kwapis et al., 2020). Here, to test the role of HDAC3, we used the selective pharmacological inhibitor RGFP966 (Bieszczad et al., 2015; Malvaez et al., 2013) to specifically block HDAC3 during the post-update reconsolidation period in 18-20-month-old mice (**FIG 1A**). During habituation, all mice showed similar decreases in movement across each session (**FIG S1A**), indicating normal habituation. We also observed low DIs during each of the three days of training, with no significant difference between groups any day of training, demonstrating there was no strong preference for object location A1 or A2 before learning (**FIG S1B;** two-tailed *t*-tests: day 1: *t*(14)=0.524, p>0.05; day 2: *t*(14)=0.771, p>0.05; day 3: *t*(14)=0.65, p>0.05).

During the update session, all mice showed intact memory for the original training information; aged mice in both groups showed a DI significantly higher than zero (**FIG 1B**; one-sample *t*-tests compared to 0: vehicle: *t*(7)=6.311, p=0.0004; RGFP966: *t*(7)=4.432, p=0.0030), and these groups were not significantly different from each other before receiving any injection (two-tailed *t*-test *t*(14)=0.4728, p=0.644). Therefore, 18-month-old mice successfully learned the original object locations with a 3-day training protocol, and all mice performed similarly before injection.

Immediately after updating, mice were given systemic injections of vehicle or RGFP966 (s.c., 10mg/kg) to block HDAC3 during reconsolidation and were tested the following day. At test, old mice given post-update RGFP966 showed normal memory for the original training locations and improved memory for the updated location compared to vehicle controls. Specifically, we found that 18-m.o. mice injected with either RGFP966 or vehicle showed intact memory for original location A1, as both groups preferentially explored novel location A4 over familiar location A1 (**FIG 1C**; one-sample *t*-tests compared to 0: vehicle: *t*(7)=3.417, p=0.0112, RGFP966: *t*(7)=3.360, p=0.0121). For location A2, as we have previously observed (Kwapis et al., 2020), old mice show weak memory, possibly due to the longer retention interval (48h) between training and testing for location A2. Although both vehicle and RGFP966 mice showed a positive DI, neither group was statistically different from 0 (one-sample *t*-test compared to 0: vehicle: *t*(7)=1.737, p=0.1260; RGFP966: *t*(7)=1.848, p=0.1071)). For both of the original locations A1 and A2, however, there were no significant group differences in DI, indicating that the groups showed similar memory for the original information regardless of drug treatment (two-tailed *t*-tests: A1: *t*(14)=1.362, p=0.195; A2: *t*(14)=0.0916, p=0.928). For the updated location A3, however, only RGFP966 mice showed intact memory, preferring to explore novel location A4 over update location A3 (**FIG 1C**; RGFP966: one-sample *t*-test compared to 0: *t*(7)=5.599, p=0.0008); vehicle-injected mice showed equal preference for locations A3 and A4 (vehicle: one-sample *t*-test compared to 0: *t*(7)=0.08333, p=0.9359), indicating no observable memory for the update, as we have previously observed in 18-m.o. mice (Kwapis et al., 2020). Further, RGFP966-injected mice had a significantly higher DI for A3 than vehicle mice (two-tailed *t*-test: *t*(14)=3.374, p=0.0045), indicating that RGFP966 enhanced memory for the updated information compared to controls. No differences in exploration time were observed between groups at test (**FIG S1C**; two-tailed *t*-test: *t*(14)=1.047, p=0.313). Together, these results demonstrate that blocking HDAC3 with RGFP966 immediately after an update can ameliorate age-related impairments in memory updating without affecting the original memory.

### Post-update HDAC3 blockade in young mice impairs the original information

Next, we wanted to see if post-update RGFP966 had any effect on young mice. As young mice successfully learn both the training and the updated information (Huff et al., 2024; Kwapis et al., 2020; Wright et al., 2020), we expected to observe no effect of post-update RGFP966. Here, young mice given one day of training were given a 5-minute memory update followed by systemic injections of RGFP966 or vehicle (**FIG 2A**). All mice were tested the following day. During habituation, all young mice showed similar decreases in movement across days (**FIG S2A**), indicating normal habituation. We also observed low DIs during training, demonstrating there was no strong preference for object location A1 or A2 before learning (**FIG S2B**; one-sample *t*-tests compared to 0: vehicle: *t*(12)=0.7250, p=0.4823; RGFP966: *t*(12)=0.6431, p=0.5323; two-tailed *t*-test comparing vehicle and RGFP966: *t*(24)=0.06; *p*>0.05).

**Figure 2.**
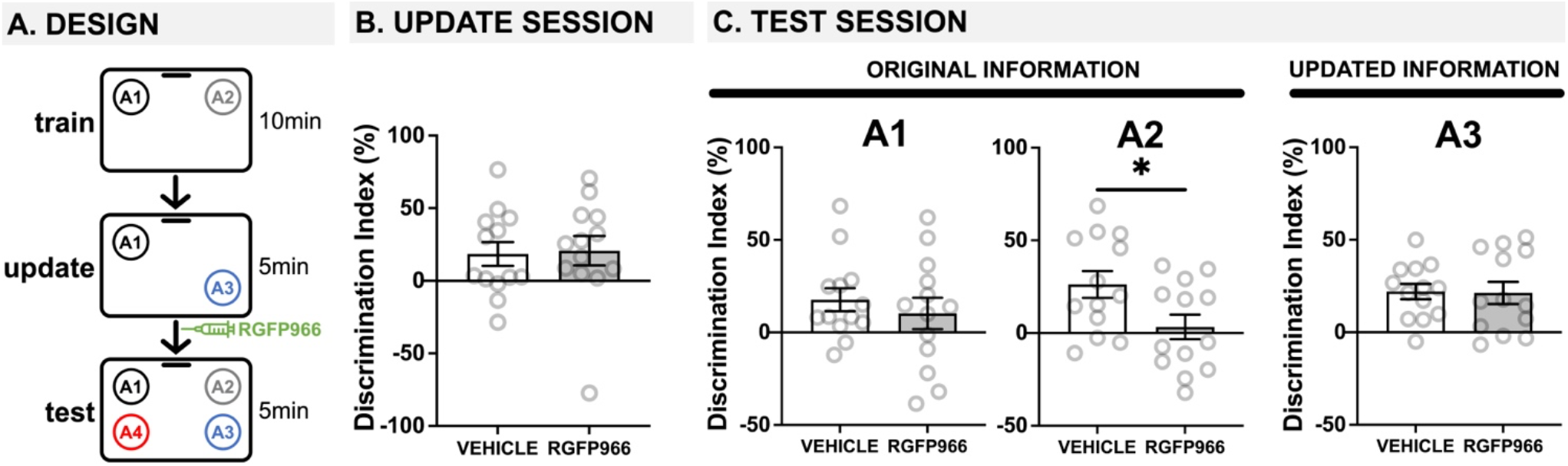
Post-update HDAC3 inhibition with RGFP966 impairs the original memory in young mice. (**A**) Experimental design using the OUL paradigm with young male mice. (**B**) Update session behavior. No differences were observed during updating, before RGFP966 injection. (**C**) Test session behavior. For locations A1 and A3, no differences were observed between RGFP966-treated and vehicle-treated animals. For the original location A2, however, post-update RGFP966 injections significantly impaired memory, with RGFP966 mice showing significantly lower DIs for A2 compared to vehicle controls. (**\***p<0.05, **\*\***p<0.01, **\*\*\***p<0.001).

During the update session, both vehicle and RGFP966 mice showed similar memory for the original training information (**Fig. 2B**; two-tailed *t*-test: *t*(24)=0.1717, *p*=0.8651). Interestingly, only animals destined to be injected with vehicle showed a DI significantly higher than zero (one-sample *t*-tests compared to 0: vehicle: *t*(12)=2.252, p=0.0438), although RGFP animals showed a positive DI that was non-significantly higher than zero (RGFP966: *t*(12)=2.042, p=0.0637). Therefore, young (3-m.o.) mice showed weak memory for the original object locations with a single training session and mice destined to receive RGFP966 or vehicle performed similarly before injection.

Immediately after the update session, mice received systemic injections of either vehicle or RGFP966 (s.c., 10mg/kg) to block HDAC3 during reconsolidation and were tested the following day. At test, young mice given post-update RGFP966 showed normal memory updating but impaired memory for the original object locations. For location A1, we found that only young mice injected with vehicle showed a DI significantly higher than zero (**FIG 2C**; one-sample *t*-tests compared to 0: vehicle: *t*(12)=2.847, p=0.0147, RGFP966: *t*(12)=1.209, p=0.2500), but we observed no significant difference between the two drug conditions (**FIG 2**; two-tailed t-test: *t*(24)=0.7085, p=0.4855). This indicates that both groups remembered the original location A1 at test, but mice injected with RGFP966 showed slightly reduced memory for the A1 location compared to vehicle controls.

Unexpectedly, however, we found that post-update RGFP966 significantly impaired memory for the original location A2. While vehicle mice showed a significant preference for location A2 at test (one-sample *t*-test compared to 0: *t*(12)=3.626, p=0.0035), RGFP966-injected mice failed to show a DI significantly higher than zero (one-sample t-test compared to 0: *t*(12)=0.4979, p=0.6276). Further, RGFP966-injected mice showed a significantly lower DI for A2 than vehicle controls (two-tailed *t*-test: *t*(24)=2.339, p=0.0280), indicating that post-update RGFP966 in young mice actually impairs memory for the original location A2. For location A3, both vehicle- and RGFP966-injected mice showed similarly intact memory; both groups had DIs significantly greater than zero (one-sample *t*-tests compared to 0: vehicle: *t*(12)=5.338, p=0.0002; RGFP966: *t*(12)=3.547, p=0.0040) and were not significantly different from each other (two-tailed *t*-test: *t*(24)=0.2080, p=0.9149), indicating successful updating. Mice injected with RGFP966 showed significantly higher total exploration than vehicle-injected mice (**FIG S2B**; two-tailed *t*-test: *t*(24)=2.509, p=0.0193), possibly reflecting this group’s impaired ability to detect that locations A2 and A1 were familiar and did not need to be explored. Therefore, in young mice that successfully update memory in OUL, blocking HDAC3 immediately after the update impairs memory for the original location A2.

### HDAC3 blockade transforms a subthreshold update into a robust memory update

We were surprised to find that blocking HDAC3 after updating actually impaired memory for the original information in young mice. One possible explanation for this effect is that the original and updated information compete for expression at test, so that in old mice, blocking HDAC3 stabilizes the weak update memory to enable its expression at test. In young mice that already successfully update, blocking HDAC3 after updating could stabilize and strengthen the memory update at the expense of the original training information, enabling the memory update to outcompete the original information so that only the updated information is properly expressed at test. To test this hypothesis, we next assessed whether blocking HDAC3 after a weak ‘subthreshold’ update would still be capable of repressing the original training information. To this end, we ran an identical experiment, except that we shortened the update session from 5 minutes to 1 minute. By reducing the update session, we created a weak update that was not capable of driving successful updating on its own in young mice, essentially mimicking the impaired updating we typically observe in old mice.

During habituation, all mice showed similar decreases in movement across each session (**FIG S3A**), indicating normal habituation. We also observed low DIs during training, with no significant differences between groups, demonstrating there was no strong preference for object location A1 or A2 before learning (**FIG S3B**; two-tailed *t*-test: t(13)=0.219, *p*>0.05). The following day, mice were all given a brief, 1-minute update session. Since this update session is so brief, mice show very low exploration times that cannot be accurately used to calculate a DI. However, young mice injected with vehicle or RGFP966 showed similar investigation times, not significantly different from each other (**FIG S3C**; two-tailed *t*-test: *t*(13)=0.235, p=0.818), suggesting similar exploration levels during the update.

As before, immediately after this weak update session, young mice were given systemic injections of vehicle or RGFP966 (s.c. 10mg/kg) and were tested the following day (FIG 3A). At test, we found that the post-update HDAC3 blockade was capable of transforming a subthreshold update into a robust update in young mice without impairing the original memory. Specifically, we found that 3-m.o. mice injected with either RGFP966 or vehicle showed intact memory for original training locations A1 and A2 (**FIG 3B**; one-sample *t*-tests compared to 0: A1 vehicle: *t*(6)=4.203, p=0.00575, A1 RGFP966: *t*(7)=4.414, p=0.0031; A2 vehicle: *t*(6)=2.666, p=0.0372, A2 RGFP966: *t*(7)=6.802, p=0.0003; two-tailed *t*-tests comparing vehicle and RGFP966: A1: t(13)=0.696, p=0.499; A2: t(13)=0.963, *p*=0.353) indicating all animals successfully recalled the original object location memory. In contrast, for the updated location A3, only RGFP966 mice showed intact memory, preferring to explore novel location A4 over update location A3 (**FIG 3B**; one-sample *t*-test compared to 0: *t*(8) = 4.395, *p*=0.0032). Vehicle mice showed equal preference for locations A3 and A4 (**FIG 3B**; one-sample t-test compared to 0: *t*(6)=0.6058, p=0.5669), indicating no observable memory for the update. This confirms that the 1-minute update was not sufficient to support memory updating in young control mice. Consistent with this, RGFP966 mice showed significantly higher DIs for location A3 than vehicle mice (**FIG 3B**; two-tailed t-test: *t*(13)=2.788, p=0.0154) with no differences in exploration time between groups (**FIG S3C**; two-tailed *t*-test: *t*(13)=1.443, p=0.173). Together, this suggests that blocking HDAC3 can strengthen a subthreshold update in young mice without impairing memory for the original training locations. Overall, this work is consistent with the idea that the original and updated information compete for expression at test, with HDAC3 inhibition shifting this balance to change which information is expressed at test.

**Figure 3.**
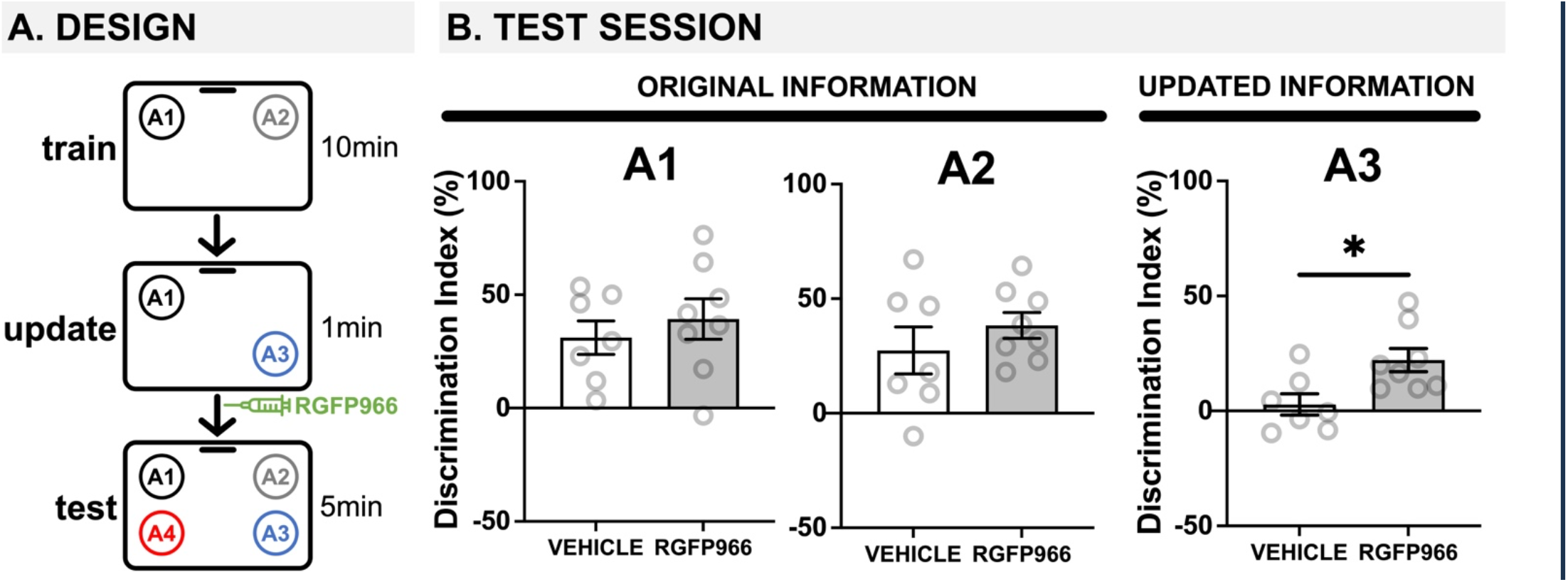
HDAC3 blockade transforms a weak update into a strong update in young mice. (**A**) Experimental design using the OUL paradigm with a subthreshold updating time of 1 minute with young male mice. (**B**) Test session behavior. No differences were observed between RGFP966-treated and vehicle-treated animals for A1 or A2, suggesting post-update RGFP966 injections did not impact memory for the original information. For A3, the subthreshold update was insufficient to drive memory updating in vehicle mice. In RGFP966 mice, however, this subthreshold update drove memory updating, with RGFP966 mice showing significantly higher DIs for A3 compared to vehicle controls. (**\***p<0.05, **\*\***p<0.01, **\*\*\***p<0.001)

## Discussion

Overall, our results show that blocking the repressive histone deacetylase HDAC3 can improve memory updating in old mice. In contrast, in young mice, we found that blocking HDAC3 after updating impairs the original memory, possibly because it creates an intensely strong update memory that can outcompete the original memory for expression at test. Together, this suggests that competition exists between the original and the updated information when a memory is updated and HDAC3 can shift the balance between which information is expressed at test. In old mice that typically show weak memory updating, RGFP966 enhances the memory update, enabling it to compete for expression with the stronger original memory. In young mice that already show robust memory updating, HDAC3 further strengthens the memory update so that it can outcompete the original information. Finally, when young mice are given a subthreshold update that alone is insufficient to support memory updating, HDAC3 can transform this weak update into one that drives robust memory updating that is expressed at test along with the original memory. Overall, this suggests that HDAC3 regulates the strength of the memory update and contributes to age-related impairments in memory updating.

Perhaps our most surprising finding was that in young mice, inhibiting HDAC3 following updating actually impaired the original memory. Enhancing the memory update by blocking HDAC3 may, therefore, come at the expense of the original training information. Only A2 was absent from the update session, suggesting that the information presented before HDAC3 inhibition becomes strengthened, whereas absent information is weakened. Inhibiting HDAC3 did not universally weaken the original information, as memory for A1 remained intact but weakened any information that was not presented during the update (A2). We observed good memory for the updated information in young animals regardless of the drug treatment. While HDAC3 inhibition did not further enhance memory for A3, the updated A3 information may have reached a ceiling for expressing behavioral enhancements in this paradigm. In the following experiments, a subthreshold update helped us to determine that this is indeed the case, as enhancing weakly updated information does not affect the original A2 information.

One interesting observation is that young mice given the HDAC3 inhibitor following a full update showed significantly more total object exploration than vehicle-treated animals, with a similar trend also observed in the subthreshold update experiment (**FIGS S2C & S3C**). While our DI calculation controls for investigation (DIs are calculated as a percentage of total exploration time), this may suggest that RGFP966-treated animals explore the objects more when they are unable to clearly detect which locations are familiar, either because the original memory was weakened by drug treatment or because the update was exceedingly brief. Indeed, movement was normal in all animals during the habituation sessions, suggesting that this difference in test exploration did not stem from an inability to move or explore normally, but rather a difference in memory for the original or updated information.

We hypothesized that competition between the original and updated information was responsible for weaker memory for the original A2 location. To test this idea, we tried weakening the update in young mice to effectively mimic the unsuccessful updating we initially observed in old mice. Specifically, we tested the effects of HDAC3 inhibition following a subthreshold update in young mice. We found that a one-minute update session was insufficient for updating unless combined with HDAC3 inhibition (**FIG 3**). Importantly, when RGFP966 was given after this subthreshold update, memory for the original information (A2) was unaffected (**FIG 3**), whereas the same inhibitor given after a full update impaired memory for the original information. This indicates that HDAC3 inhibition can manipulate the balance between the original and updated information; blocking HDAC3 strengthened the update information, creating either an intensely robust memory that outcompeted the original information or transforming a weak update into one capable of competing for expression with the original memory.

Our work, therefore, indicates that the repressive histone deacetylase HDAC3 plays a key role in reconsolidation-based memory updating, and inhibiting HDAC3 can improve memory updating in old mice. This is consistent with the well-documented role of HDAC3 in regulating memory strength and persistence across paradigms (Alaghband et al., 2017; Campbell et al., 2021; Hervera et al., 2019; Hitchcock et al., 2019; Keiser et al., 2021; Kwapis, Alaghband, et al., 2017; Kwapis et al., 2019; Malvaez et al., 2013; McQuown et al., 2011; McQuown & Wood, 2011; Rogge et al., 2013; Shu et al., 2018). This is a key finding, considering the lack of information about the specific roles of epigenetic mechanisms in memory reconsolidation or memory updating. Past work using less selective HAT or HDAC inhibitors has demonstrated impairment or enhancement of reconsolidation in fear conditioning, respectively, with an emerging role for Class I HDACs (including HDACs 1, 2, 3, and 8; Bredy & Barad, 2008; Gräff et al., 2014; Maddox, Watts, Doyère, et al., 2013; Maddox, Watts, & Schafe, 2013; Maddox & Schafe, 2011). Inhibitors specific to Class I HDACs (including VPA and Cl-994) are sufficient to improve reconsolidation (Bredy & Barad, 2008; Gräff et al., 2014). Here, our work indicates that HDAC3 plays a crucial role in modulating the strength of reconsolidation-dependent memory updating in both young and old mice. In addition to work showing a key role for DNA methyltransferase activity in reconsolidation (Maddox et al., 2014; Maddox, Watts, & Schafe, 2013), it is becoming increasingly clear that a range of epigenetic mechanisms may be critical for successful memory reconsolidation and, possibly, memory updating.

Although the OUL memory updating paradigm is new, a number of labs have used this task to better understand how memory updating works. Previous work has confirmed that OUL drives memory updating rather than the formation of a new memory. First, blocking protein synthesis within the dorsal hippocampus after the update session impairs memory for both the original training information and the update, suggesting the original memory is destabilized in response to the updated location (Kwapis et al., 2020). Second, using *Arc* catFISH, we have previously confirmed that the update session preferentially reactivates the neuronal ensemble supporting the original object location memory (Kwapis et al., 2020). OUL, therefore, appears to drive reconsolidation-based updating in the original neuronal ensemble. Recent work has demonstrated that muscarinic acetylcholine receptors (mAChRs) are required for the original memory to destabilize during updating; blocking mAChRs with scopolamine before updating prevented mice from learning the updated location A3 without affecting memory for the original locations A1 and A2 (Huff et al., 2024). Finally, exposing mice to galactic cosmic radiation (intended to mimic space radiation for astronauts traveling to Mars) disrupts memory updating even when initial memory formation is intact (Alaghband et al., 2023; Keiser et al., 2021). Although this work is in the early stages, the OUL paradigm is poised to support a better understanding of the mechanisms that support memory destabilization, restabilization, and information integration at the molecular, cellular, and behavioral levels.

The current experiments show evidence for HDAC3-mediated memory competition among original and updated object location memories. To better understand what HDAC3 is doing during memory updating, future studies should focus on identifying which genes are mediated by HDAC3 during updating and determine if these genes are identical to or different from the genes required for initial memory consolidation. Determining how these transcriptional profiles change in old age when memory updating is impaired will also be critical. Finally, the current experiments were all done in male mice, and subsequent studies will need to add female mice to determine if these mechanisms are similar in both sexes. While our lab has just started to use the OUL paradigm with female mice, the molecular mechanisms by which memories compete may differ in females.

Overall, our findings show that HDAC3 regulates the competition between an original memory and a memory update and suggest that HDAC3 may contribute to age-related impairments in memory updating. Our findings demonstrate that epigenetic mechanisms play a key role in memory updating and suggest that OUL is poised to help identify the molecular, cellular, and circuit-level mechanisms that support hippocampal memory updating in rodents.

## Author Contributions

JLK and CWS designed the experiments. JLK, CWS, LB, DSW, SB, MWU, CAB, and GS collected the data. JLK, CWS, and GS analyzed the data and wrote the manuscript. CWS prepared the figures.

## Supplementary Material

*Supplementary materials have been provided in an attached document*.

