## Supplemental Material for "Pharmacological HDAC3 inhibition alters memory updating in young and old mice"

### *Supplementary Material*

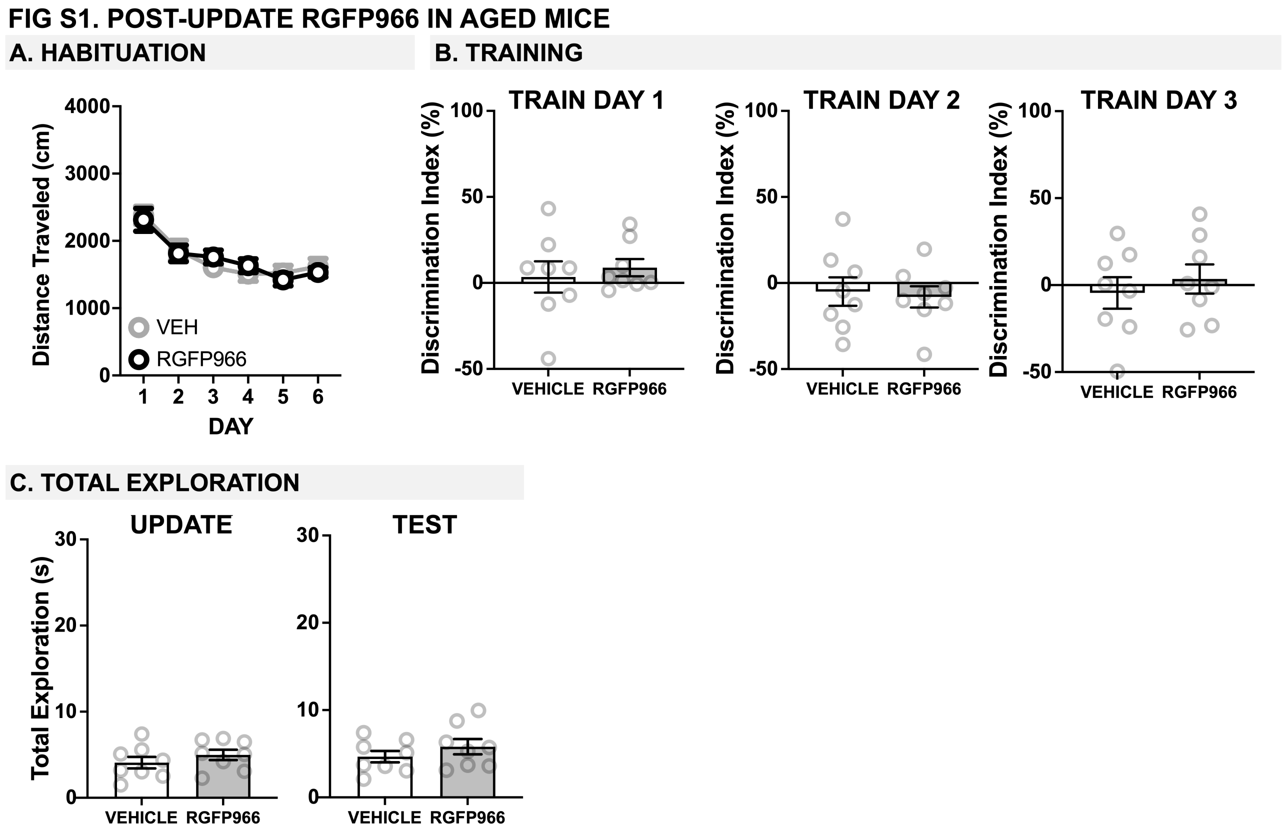


**Supplementary Figure 1.** *Post-update RGFP966 in aged mice* (**A**) Mice decreased their movement over six days during habituation, with similar distance traveled for vehicle and RGFP966 mice before injection. (**B**) Training DIs for each day of training. Vehicle and RGFP966 mice show similar, low DIs across each day. **(C)** Total exploration during the test session was similar for vehicle and RGFP966 mice. (*p<0.05, **p<0.01, ***p<0.001)

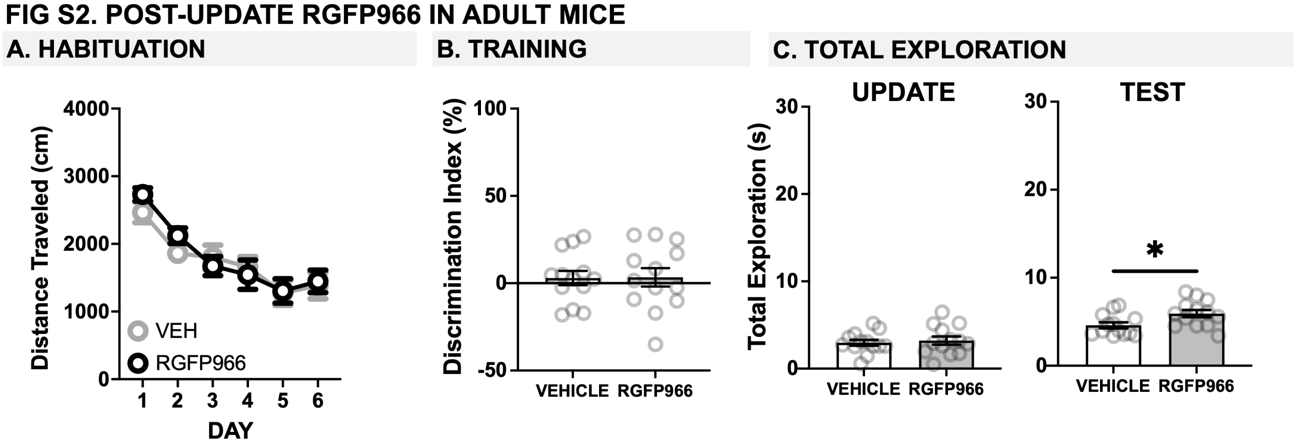


**Supplementary Figure 2.** *Post-update RGFP966 in young mice* (**A**) Mice decreased their movement over six days during habituation, with similar distance traveled for vehicle and RGFP966 mice before injection. (**B**) Training DIs for training. **(C)** Total exploration during the test session was higher for RGFP966 compared to vehicle mice. (*p<0.05, **p<0.01, ***p<0.001)

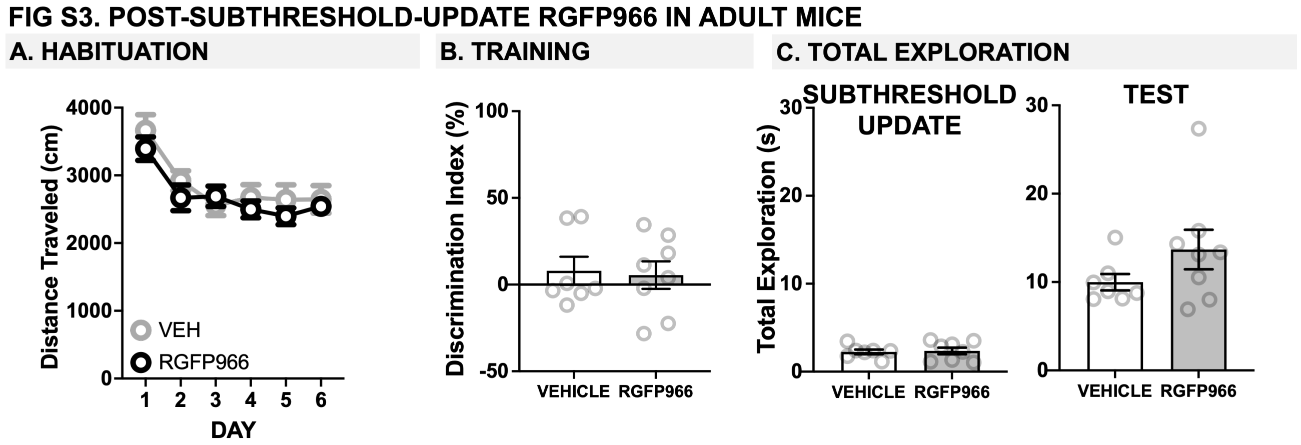


**Supplementary Figure 3.** *Post-subthreshold-update RGFP966 in adult mice* (**A**) Mice decreased their movement over six days of habituation, with similar distance traveled for vehicle and RGFP966 mice before injection. (**B**) Training DIs were similar for vehicle and RGFP966 mice before injection. **(C)** Total exploration during the test was similar for vehicle and RGFP966 mice. (* $p<0.05$, ** $p<0.01$, *** $p<0.001$)
